# Screening of different cytotoxicity methods for the assessment of ENDS toxicity relative tobacco cigarettes

**DOI:** 10.1101/2021.02.25.432848

**Authors:** Massimo Caruso, Rosalia Emma, Sonja Rust, Alfio Distefano, Giuseppe Carota, Roberta Pulvirenti, Riccardo Polosa, Giovanni Li Volti

## Abstract

Electronic Nicotine Delivery Systems (ENDS), i.e., electronic cigarettes (e-cigs) and Tobacco Heating Products (THPs), are rapidly growing in popularity. The marketing of these products is regulated by specific rules in the European Union and in the US, which permit their legal sales. Nonetheless, comprehensive quality and safety requirements for regulatory purposes are still under development. Cytotoxicity studies are an important initial step in appraising the potential toxicity of ENDS. The aim of the present study was to screen a battery of different in vitro cytotoxicity methods for the assessment of toxicity induced by ENDS. We evaluated different cytotoxicity assays, including neutral red uptake (NRU), 3-(4,5-dimethylthiazol-2-yl)-2,5-diphenyltetrazolium bromide (MTT), Annexin V apoptosis, High Content Screening (HCS) assays and Real Time Cell Analysis (RTCA), to compare two e-cigs (Vype ePen 3 and Vype eStick Maxx) and two THPs (IQOS and GLO™) with the 1R6F reference tobacco cigarette. Human bronchial epithelial cells (H292) were exposed to 1R6F smoke (5 puffs by HCI regime), ePen vapor (10 puffs by modified HCI regime), eStick vapor (25 puffs by CRM81 regime), IQOS vapor (7 puffs by HCI regime) and GLO vapor (8 puffs by HCI regime) at air-liquid interface. All tests showed reduced cell viability following 1R6F smoke exposure and slight or no reduction with ENDS at 24 hours compared to controls. In addition, Annexin V and RTCA exhibited a further significant reduction in cell viability following 1R6F exposure compared with other assays. Furthermore, Annexin V allowed to discriminate viable cells from those in early/late apoptosis. Finally, RTCA and HCS being time-resolved analyses allowed also to determine the kinetic dependency parameter for toxicity of smoke/vapor chemicals on cell viability. In conclusion, NRU assay may be considered a suitable test, especially when combined with a time-resolved test, for assessing the kinetic of cytotoxicity induced by these products.

## 1. Introduction

Cytotoxicity assays have been widely used to assess the toxicological impact of tobacco smoke (Belushkin et al., 2014). Indeed, regulatory authorities included *in vitro* toxicity tests in a battery of assays to detect all possible toxicity effects of tobacco products (Belushkin et al., 2014; CORESTA, 2004). Consequently, these toxicity assays are routinely used as standard methods by tobacco industry for product assessments (Baker et al., 2004a, 2004b; Dempsey et al., 2011). More recently marketing of alternative tobacco products, generally referred as electronic nicotine delivery systems (ENDS), including electronic cigarettes (e-cig) and tobacco heating products (THPs), increased the need to properly assess their potential toxic effects (Johnson et al., 2009). Currently, alternative tobacco products are tested following the standard approach used for tobacco products, but a more specific approach for e-cig and THPs could improve their toxicity assessment (Iskandar et al., 2016). However, the rapid evolution of these products, and their large variability, due to the lack of manufacturing standard, make it difficult to develop standard protocols for toxicity assessment (Davis et al., 2015; Iskandar et al., 2016; Kim et al., 2015). Neutral red uptake (NRU) is the most used assay for cytotoxicity evaluation in the context of tobacco products testing (Belushkin et al., 2014), and it is also included in the first non-genotoxicity *in vitro* assay accepted for the regulatory evaluation of chemical compounds (European Commission, 2000; OECD/OCDE, 2004; Repetto et al., 2008). The NRU cytotoxicity test is a cell survival/viability chemosensitivity test based on the ability of viable cells to incorporate and bind neutral red (NR), a weak supra-vital cationic dye that freely penetrates cell membranes by non-ionic diffusion and accumulates in lysosomes. Alterations of biological membranes leading to lysosomal fragility and other changes caused by the action of the chemical mixture (e.g., cigarette smoke) can result in a decrease in the absorption and binding of NR. This makes it possible to distinguish between viable, damaged or dead cells. The degree of inhibition of growth, related to the concentration of the test compound, provides an indication of cytotoxicity expressed as a reduction in NR absorption after chemical exposure, thus providing a sensitive signal of both cell integrity and growth inhibition (Putnam et al., 2002; Repetto et al., 2008; The National Toxicology Program (NTP) Interagency Center for the Evaluation of Alternative Toxicological Methods (NICEATM), 2003). Other commonly used methods for cytotoxicity evaluation include measurement of the reduction of tetrazolium salts. The yellow 3-(4,5-dimethyl-2-thiazolyl)-2,5-diphenyl-2H-tetrazolium bromide (MTT, thiazolyl blue) is the most used among them. The water-soluble MTT salts are incorporated by viable cells due to their net positive charge, and then reduced by dehydrogenases and other reducing agents contained in metabolically active cells (Berridge et al., 2005; Stockert et al., 2018). The reduction of MTT leads to the formation of insoluble violet-blue formazan product, which are proportional to cell viability (Berridge et al., 2005; Johnson et al., 2009). The use of the above-described methods has some drawbacks since they do not discriminate among apoptosis, necrosis or autophagy thus leading to a possible underestimation of toxicity. Indeed, the induction of programmed cell death (i.e., apoptosis) and necrosis processes represent crucial steps in the evaluation of cytotoxicity. In particular, though apoptosis is not an inflammatory form of programmed cell death, it is well known that cigarette smoke induces apoptosis in human bronchial epithelial cells (Comer et al., 2013; Imai et al., 2005). Annexin V evaluation by cytofluorimetric analysis to assess apoptosis represents a useful tool to measure of cytotoxicity even though it requires specific laboratory skills and instruments.

Recently, high-throughput technology has made possible to detect cellular changes with an overall overview of biological response and monitoring them continuously over time (Iskandar et al., 2016). These new rapid methods, alongside with the classic cytotoxicity assay, could provide a deeper knowledge on toxicity of alternative tobacco products. Two of these new methods are the High content screening (HCS)-based multiparametric analysis and the Real-Time Cell-based Assay (RTCA) technology. HCS analysis represents a useful tool in early toxicity testing because it allows to measure simultaneously numerous parameters including mitochondrial membrane potential, plasma membrane permeability, oxidative stress and morphological parameters without the use of specific dyes (Mandavilli et al., 2018). Instead RTCA, a real-time cellular biosensor, allows for uninterrupted, label free, and real time analysis of cells over the course of an experiment with the advantages of avoiding marks, violation to the cell, and overcoming the interference of the compounds in detection (Yan et al., 2018).

Since there is not a specific indication or protocol for toxicity evaluation of e-cig and THPs products on human bronchial epithelial cells, we evaluated a number of *in vitro* cytotoxicity assays (i.e., NRU assay, MTT assay, Annexin V apoptosis assay, NRU assay, HCS and RTCA technologies) by comparing two brands of e-cig (Vype ePen 3 and Vype eStick Maxx), two THPs (IQOS and GLO™) with the 1R6F research cigarettes using human bronchial epithelial cells exposed to smoke/vapor at air-liquid interface (ALI).

## 2. Methods

### 2.1 Test products and exposure regimes

1R6F reference cigarettes (Center for Tobacco Reference Products, University of Kentucky) were used for smoke exposure. These cigarettes have been reported to produce 46.8 mg TPM, 29.1 mg tar and 1.896 mg nicotine per cigarette smoked following HCI regime (Center for Tobacco Reference Products Kentucky University, 2018). Cigarettes were conditioned at 22±1 °C and 60±3 % of relative humidity for at least 48 hours according to ISO 3402:1999 guidelines. LM1 smoking machine (Borgwaldt KC GmbH, Hamburg – Germany) was used to smoke 5 puffs of 1R6F cigarette following HCI regime which ensures a 55 ml, 2 s duration bell shape profile, puff every 30 s with filter vent blocked. Vapor exposure was carried out using two electronic cigarettes, Vype ePen 3 and Vype eStick (British American Tobacco; http://www.govype.com) (Azzopardi et al., 2016) and two THPs, IQOS Duo (Philip Morris Products SA; https://it.iqos.com) and GLO™ Pro (British American Tobacco; https://www.discoverglo.com). Vype ePen3 is a closed-system e-cigarette with a cotton wick comprising a rechargeable 650 mAh battery and actuation button, and a disposable cartridge with integral mouthpiece. The device has a single 6 W power setting. Vype eStick is puff-activated cigarette-like product consisting of two modules, a rechargeable battery section and a replaceable liquid (“e-liquid”) containing cartridge (“cartomizer”) operating at 3.7 V (Azzopardi et al., 2016). IQOS Duo consists of three distinct components that perform different functions: (i) a tobacco stick containing a tobacco powder with the addition of water, glycerin, guar gum and cellulose fibers, (ii) a holder into which the tobacco stick is inserted and which heats the tobacco material by means of an electronically controlled heating blade from inside the tobacco stick, and (iii) a charger that is used to recharge the holder after one or two uses. The temperature of the heating blade is carefully controlled and the operating temperature does not exceed 350°C (Bekki et al., 2017; Li et al., 2019). GLO™ Pro includes two parts: an electronic handheld device with a heating chamber equipped with a rechargeable Li-ion battery (3000 mAh capacity), and a custom-made tobacco rod to be inserted into the heating chamber. The electronic device has a heating module separately controlled by the inbuilt software, and thus, the tobacco rod is heated to less than 250°C from the periphery. This is significantly lower than the major pyrolysis and combustion temperature ranges seen in a lit cigarette (typically between 350°C and 900°C) but is sufficient to release nicotine, glycerol (added as the main aerosol agent) and volatile tobacco flavor compounds (Eaton et al., 2018). “Master Blend” flavored variant containing 18 mg/mL nicotine was used for Vype ePen3, “Toasted Tobacco” flavored variant containing 18 mg/mL nicotine was used for Vype eStick Maxx, Heets “Sienna Selection” was used for IQOS duo, and Neo™ Sticks “Ultramarine” was used for GLO™ Pro. Vapor exposure was performed using LM4E vaping machine (Borgwaldt KC GmbH, Hamburg – Germany). Vype ePen3 (button-activated e-cigarette) was vaped following a modified HCI regimen (55 mL puff volume, drawn over 2 s, once every 30 s with square shape profile) plus 1 s of pre-activation, for 10 puffs. Vype eStick Maxx was vaped following CRM81 regimen (55 mL puff volume, drawn over 3 s, once every 30 s with square shape profile) for 25 puffs. IQOS Duo and GLO™ Pro were vaped using HCI regime without blocking the filter vents, to avoid the device overheating, for 7 and 8 puffs respectively. The ENDS product batteries were fully charged before use and a fresh e-liquid cartridge for e-cig were used. The puff number for each product was established according to nicotine dose delivered from 1R6F causing the 50% of cell death (data not shown), in order to have the similar nicotine delivery for the other test products.

### 2.2 Cell culture and air-liquid interface (ALI) exposure methods

Human adenocarcinoma lung epithelial cells (NCI-H292, ATCC® CRL-1848™) were cultured as described previously by Azzopardi et al. 2015 (1). Briefly, H292 cells were cultured in RPMI 1640 medium (10% foetal bovine serum, 2 mM L-glutamine, 50 U/ml penicillin and 50 mg/ml streptomycin) at 37 °C, 5% CO2 in a humidified atmosphere. Then, cells were seeded in 12 mm Transwells® inserts (Corning Incorporated, NY, USA) at a density of 3×10^5^ cells/ml sustained by 1 ml of RPMI medium in the basal compartment of each well and 0.5 ml in the apical compartment of each Transwell® insert, 48 hours prior to exposure. Cell starvation was done 24 hours prior to exposure by replacing the basal and apical medium with 1 mL and 0.5 mL respectively of UltraCULTURE™ containing 2 mM glutamine, 50 U/mL penicillin and 50 μg/mL streptomycin. Next, when the 80% confluency was reached, the apical medium was removed from each insert and two inserts per test product were transitioned to the exposure chamber with 20 ml of DMEM-high glucose (DMEM-hg) in the basal compartment in order to perform the air-liquid interface (ALI) exposure (Fig. 1). This exposure method is the most physiologically relevant for bronchial epithelial cell lines (e.g., H292) exposing them to all fractions and components of smoke/vapor (Azzopardi et al., 2015). For each smoking/vaping regime, one exposure chamber was connected to the LM4E port without the device so as to expose H292 cells to laboratory air filtered by a Cambridge Filter Pad at the same regime (AIR control). Moreover, 2 negative controls, consisting of 1 seeded insert with media submerged (INC) and 1 seeded insert without apical media (ALI) in the incubator, and 1 positive control with 1 ml apical and 2 ml basal sodium dodecyl sulphate (SDS) at 350 μM were included for each set of exposure. After each exposure, the inserts were transferred from the chamber to a clean well plate, adding 1 mL and 0.5 mL of supplemented UltraCULTURE™ respectively at the basal and apical side for 24 hours of recovery period. The recovery period was not performed for Neutral Red Uptake (NRU) Assay in live and xCELLigence Real-Time Cell analysis.

**Figure 1.**
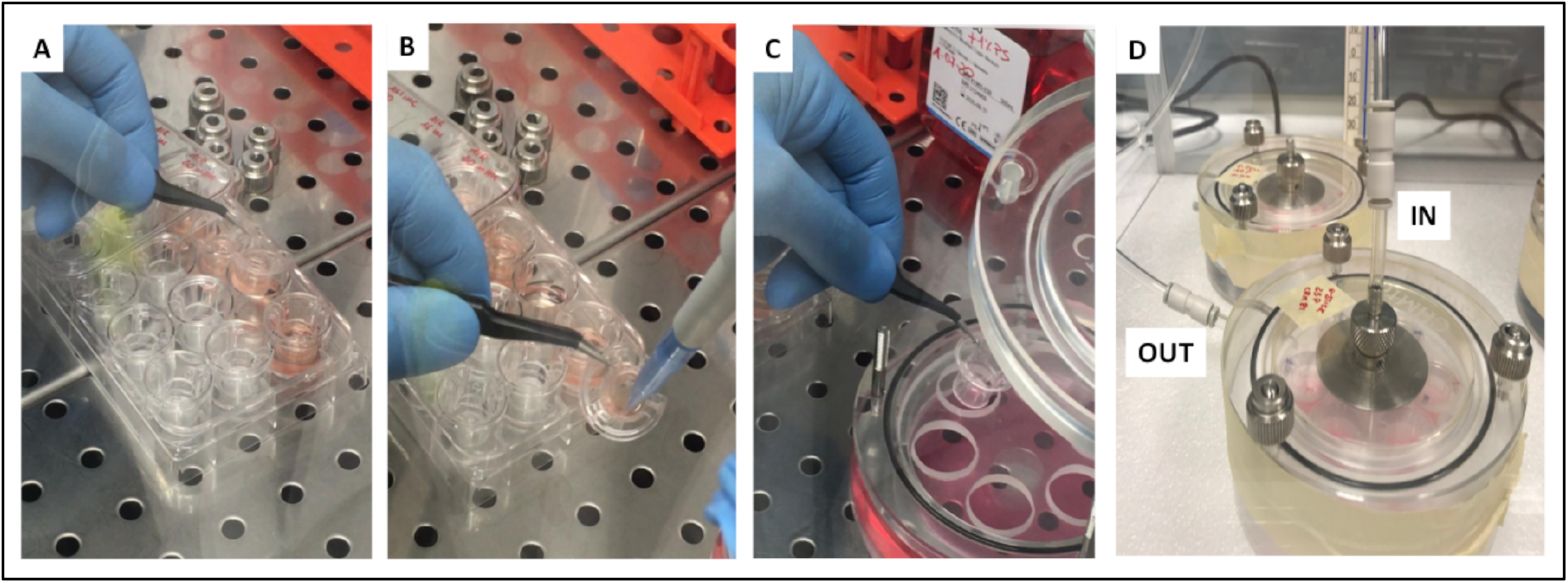
Exposure at AIR-Liquid Interface (ALI). (A) H292 cells are seeded in Transwell® inserts. (B) At 80% confluency, the apical medium is removed from the inserts, and then (C) transitioned to the exposure chamber with 20 ml of DMEM-hg in the basal compartment. (D) The exposure chamber is placed in a controlled temperature incubator (37°C) and connected with smoking/vaping machine through the IN tube. An OUT tube is connected to the exhaust.

### 2.3 Neutral Red Uptake (NRU) Assay

After 24 h recovery period, UltraCULTURE™ medium was removed and exposed cells were washed twice with phosphate buffered saline (PBS). Then, cells were incubated with Neutral Red (NR) dye (0.05 g/L in UltraCULTURE™) for 3 h at 37 °C, 5% CO^2^ in a humidified atmosphere. Next, other two washes with PBS were done to remove unincorporated dye. 500 μl of destain solution (50% ethanol, 49% distilled water, 1% glacial acetic acid; V:V:V) was added to each insert in order to elute incorporated NR from cells by incubation for 10 min at 300 rpm on a plate shaker. NR extracts were transferred to a 96-well plate in triplicate, in aliquots of 100 μl per well. The optical density of NR extracts was read with a microplate spectrophotometer (Synergy HT, BioTek) at 540 nm using a reference filter of 630 nm. A blank insert (without cells) was used to assess how much NR solution stains the Transwells® membranes. Background measurement from Blank was subtracted from each measurement. NRU levels of treated cells were expressed as a percentage of air-exposed controls.

### 2.4 MTT Assay

Basal and apical UltraCULTURE™ medium was removed after 24 hours of recovery. Cells were washed twice with PBS and then incubated with 1.5 ml (0.5 ml apical and 1 ml basal) of 0.5% 3-(4,5-dimethylthiazol-2-yl)-2,5-diphenyltetrazolium bromide (MTT) in UltraCULTURE™ for 3 hours at 37 °C, 5% CO^2^ in a humidified atmosphere. Next, 500 μl of DMSO was added to each insert and incubated for 10 min at 300 rpm on a plate shaker in order to dissolve the formazan crystals produced. The eluted formazan crystals were transferred to a 96-well plate in triplicate, in aliquots of 100 μl per well. The optical density of MTT extracts was read with a microplate spectrophotometer (Synergy HT, BioTek) at 570 nm. A blank insert (without cells) was used to assess how much MTT salts were trapped in the Transwells® membranes. Background measurement from Blank was subtracted from each measurement. Cell viability was expressed as percentage of air-exposed controls.

### 2.5 Annexin V Apoptosis Assay

Evaluation of apoptosis and necrosis was performed using the Muse® Annexin V & dead cell Kit (Luminex Corporation, Austin - USA). After a recovery period of 24 hours, H292 cells were washed, trypsinized (0.25% trypsin) and resuspended in supplemented RPMI-1640 medium. Each exposure condition was assayed in duplicate. The assay was carried out following the manufacturers’ instructions. Viable cells [Annexin V-PE (−) and 7AAD (−)], early apoptotic cells [Annexin V-PE (+) and 7AAD (−)], advanced apoptotic cells [Annexin V-PE (+) and 7AAD (+)], and dead cells [Annexin V-PE (−) and 7AAD (+)] were evaluated as percentage gated. The percentage of viable cells was expressed as percentage of HCI-AIR control when compared with the results of other assays.

### 2.6 Neutral Red Uptake (NRU) Assay in live imaging

Exposed H292 cells were washed twice with PBS and detached from the inserts with 0.25% trypsin. Cells were seeded in a CellCarrier™-96 well (PerkinElmer) at a density of 1×10^4^ cells/ml, and incubated at 37 °C, 5% CO^2^ for 24 hours. Next, the cells were labelled with 100 μl of a solution consist of 0.05 g/L NR dye and 2 droplets/ml NucBlue™ (Invitrogen) in UltraCULTURE™ by incubating for 3 h at 37 °C, 5% CO^2^ in a humidified atmosphere. After the 3 hours of staining, medium was removed from each well and cells were washed twice with PBS. Then, 200 μl of fresh supplemented RPMI-1640 medium was added in each well. The plate was read under confocal conditions using the 20x long WD objective by High Content Screening (HCS) analysis (PerkinElmer Operetta High-Content Imaging System). Exposed H292 cells were monitored every 1 h until 24 h and, then every 4 h until 48 h. All images were analysed using Harmony high-content imaging and analysis software (PerkinElmer). Final output values from the analysis are expressed as mean fluorescence intensity (MFI) percentage of control per well. Live cell viability curves were generated for each tested product.

### 2.7 xCELLigence Real-Time Cell analysis

At the end of each exposure, H292 cells were washed twice with PBS, trypsinized (0.25% trypsin), counted and resuspended in supplemented RPMI-1640. Next, cells were seeded in E-16 xCELLigence plate at a density of 3 x 10^3^ cells/ml per well. The plates were then incubated at 37 °C, 5% CO^2^ for 30 min in order to allow cell settling. Real-time cell proliferation analysis was performed using the xCELLigence RTCA DPsystem. Real-time changes in electrical impedance were measured and expressed as “*cell index*”, defined as (Rn-Rb)/15, where Rb is the background impedance and Rn is the impedance of the well with cells. The background impedance was measured in E-plate 16 with 100μL medium (without cells) after 30 min incubation period at room temperature. Cell proliferation was monitored every 20 min for 72 hours.

### 2.8 Statistics

Distribution of data was assessed using the Shapiro-Wilk test. Data were summarised using the mean±standard deviation (SD). Comparisons of NRU, MTT, Annexin V Apoptosis results, and comparisons among the assays were made using one-way analysis of variance (ANOVA) with differences between groups determined using Tukey’s adjustment for multiple comparisons. For NRU in live imaging and xCELLigence Real-Time Cell analysis, p values were calculated by applying two-way ANOVA with differences between groups determined using Tukey’s adjustment for multiple comparisons. Moreover, linear regression and multiple linear regression analyses were performed in order to verify the relationships among the cell viability curves. All analyses were considered significant with a p-value of less than 5 %. Analyses of data were performed using R version 3.6.1 (R Core Team, 2019).

## 3. Results

### 3.1 Cell viability by NRU assay

NRU cell viability assay showed that all used products had a significant difference with an overall p value< 0.0001 (Fig. 2). Particularly, we observed a significant reduction of cell viability after exposure to 1R6F smoke (32.02% ± 1.78) compared to AIR-HCI control (p< 0.0001). Similarly, significant cell viability decrease was observed after exposure to IQOS (88.35% ± 3.88) (p< 0.001), and GLO (89.98% ± 3.99) (p< 0.01) compared to AIR-HCI control. On the other hand, H292 exposure to ePen vapor showed no reduction in cell viability (95.63% ± 3.97) compared to AIR-mHCI control (p> 0.05). No reduction in cell viability was also observed in H292 cells exposed to eStick vapor (101.59% ± 4.06) compared to AIR-CRM81 control. Cross comparisons among the study products showed that cell viability reduction after 1R6F exposure is significantly different when compared to all the other products (p values < 0.0001). Furthermore, significant differences were observed for IQOS *vs* eStick (p< 0.0001), IQOS *vs* ePen (p= 0.006), GLO *vs* eStick (p< 0.0001), and ePen vs eStick (p= 0.042). Instead, no significant differences were observed for GLO *vs* ePen (p= 0.063) and for GLO vs IQOS (p= 0.981).

**Figure 2.**
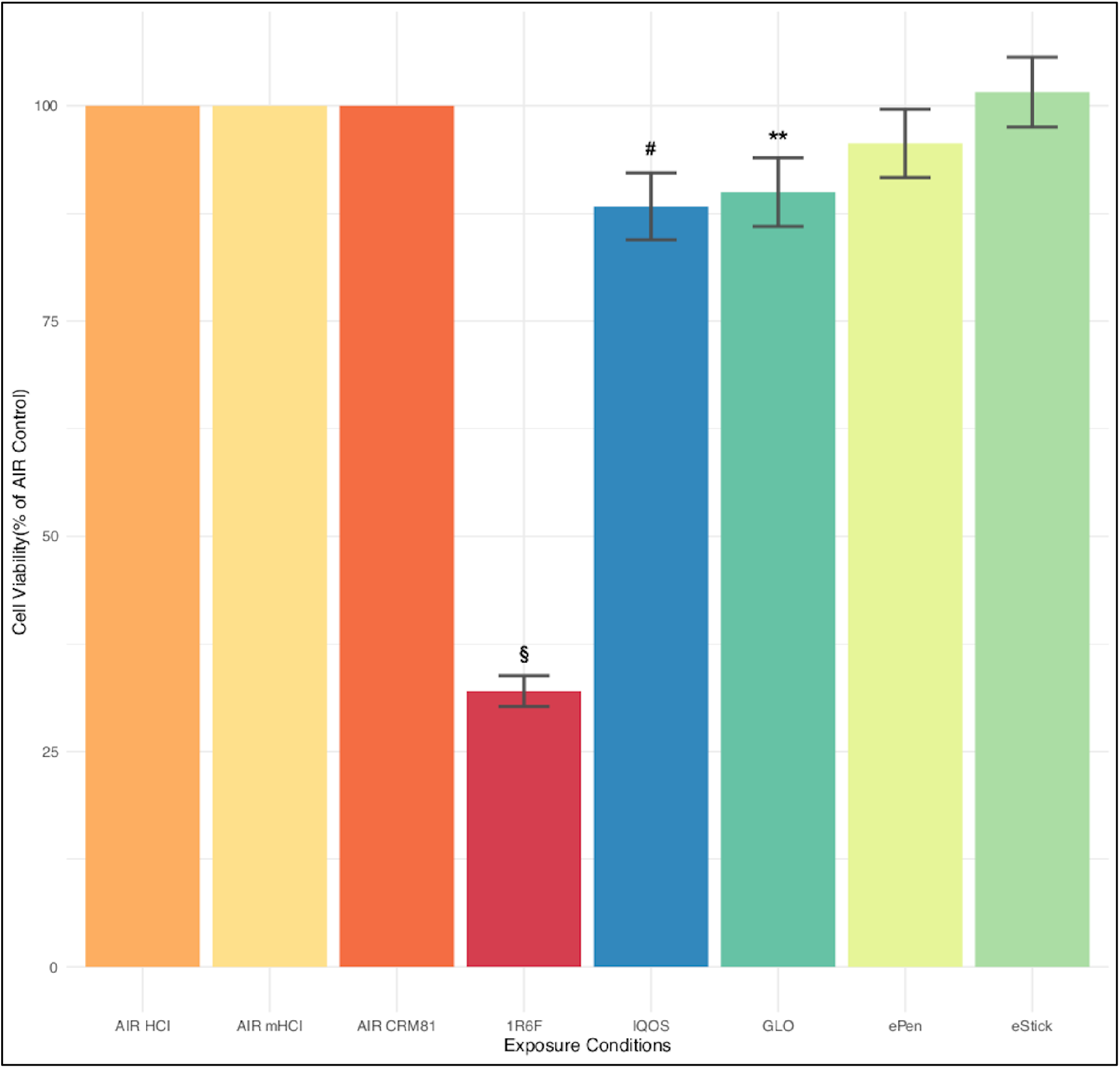
Evaluation of cell viability by NRU assay. Cell viability of each tested product is expressed as percentage of its AIR control. The mean ± SD values were respectively, 32.02 ± 1.78% for 1R6F, 88.35 ± 3.88% for IQOS, 89.98 ± 3.99% for GLO, 95.63 ± 3.97% for ePen, and 101.59 ± 4.06% for eStick. Significance code: p< 0.0001 (§) *vs* AIR control; p< 0.001 (#) *vs* AIR control; p< 0.01(**) *vs* AIR control.

### 3.2 Cell viability by MTT assay

Consistently with NRU assay we observed a significant difference among the products tested in this study with an overall p value< 0.0001 (Fig. 3). Considerable H292 cell viability reduction was shown after exposure to 1R6F smoke with 49.21 ± 9.95% of viable cells, compared to AIR-HCI control (p< 0.0001). Instead, the means±SD of cell viability percentage after exposure to IQOS, 97.29 ± 9.95 %, GLO, 96.79 ± 8.54 %, ePen, 90.08 ± 2.79 %, and eStick, 86.69 ± 3.63 %, were not different from their related AIR controls. Moreover, cross comparisons among these products showed no significant differences in cell viability percentages (p values> 0.05). Conversely, cell viability after exposure to IQOS, GLO, ePen, and eStick was significantly higher compared to cell viability after 1R6F smoke exposure (p values < 0.0001).

**Figure 3.**
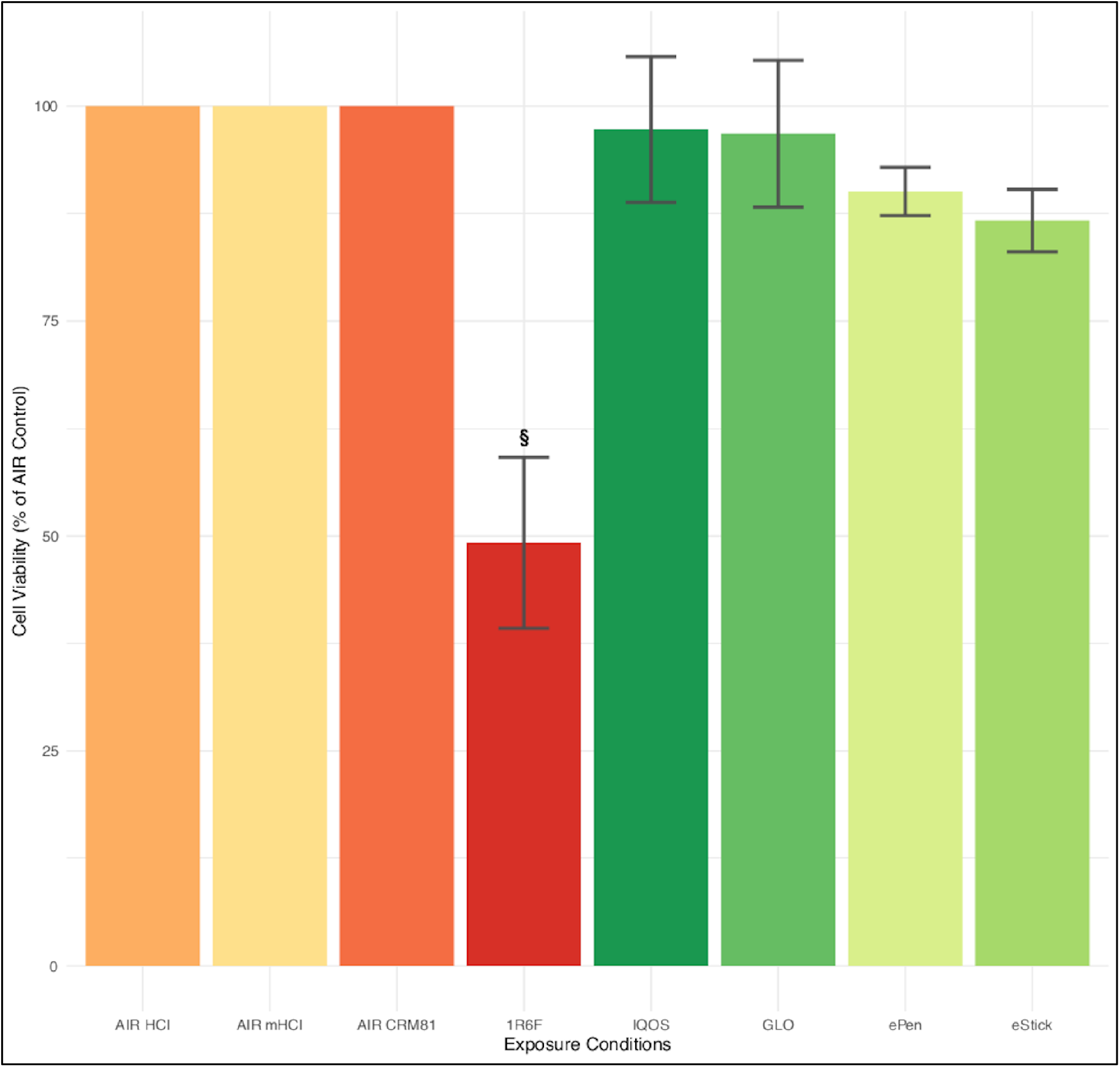
Evaluation of cell viability by MTT assay. Cell viability of each tested product is expressed as percentage of its AIR control. The mean ± SD values were respectively, 49.21 ± 9.95% for 1R6F, 97.29 ± 9.95 % for IQOS, 96.79 ± 8.54 % for GLO, 90.08 ± 2.79 % for ePen, and 86.69 ± 3.63 % for eStick. Significance code: p< 0.0001 (§) *vs* AIR control.

### 3.3 Evaluation of apoptosis and necrosis after smoke/vapor exposure

Flow cytometric evaluation of apoptosis and necrosis of H292 cells exposed to 1R6F cigarette smoke, THPs (IQOS and GLO), and e-cig (ePen and eStick) is shown in Figure 4. The mean ± SD of viable cells percentage gated is significantly lower for cells exposed to 5 puff of 1R6F cigarette, 7.35 ± 1.2 %, compared to exposure with IQOS, 66.95 ± 6.7%, GLO, 68.55 ± 0.91%, ePen, 69.55 ± 0.4%, eStick, 73.55 ± 0.8%, with p values <0.0001. Moreover, significant increase of advanced apoptotic cell percentage was observed for cells exposed to 1R6F cigarette, 67.15 ± 2.6 %, compared to cells exposed to IQOS, 10.2 ± 2.8%, GLO, 10.15 ± 0.2%, ePen, 8.9 ± 0.6%, eStick, 7.5 ± 1%, (p values <0.0001). Also, the percentage of early apoptotic cells after exposure to 1R6F cigarette, 23.55 ± 1.5%, was significantly different from AIR-HCI control, 15.75 ± 1.2% (p= 0.01), and eStick, 16.6 ± 1.7%, (p= 0.02). No differences were observed between early apoptotic cells after exposure to 1R6F cigarette and early apoptotic cells after exposure to IQOS, GLO, and ePen.

**Figure 4.**
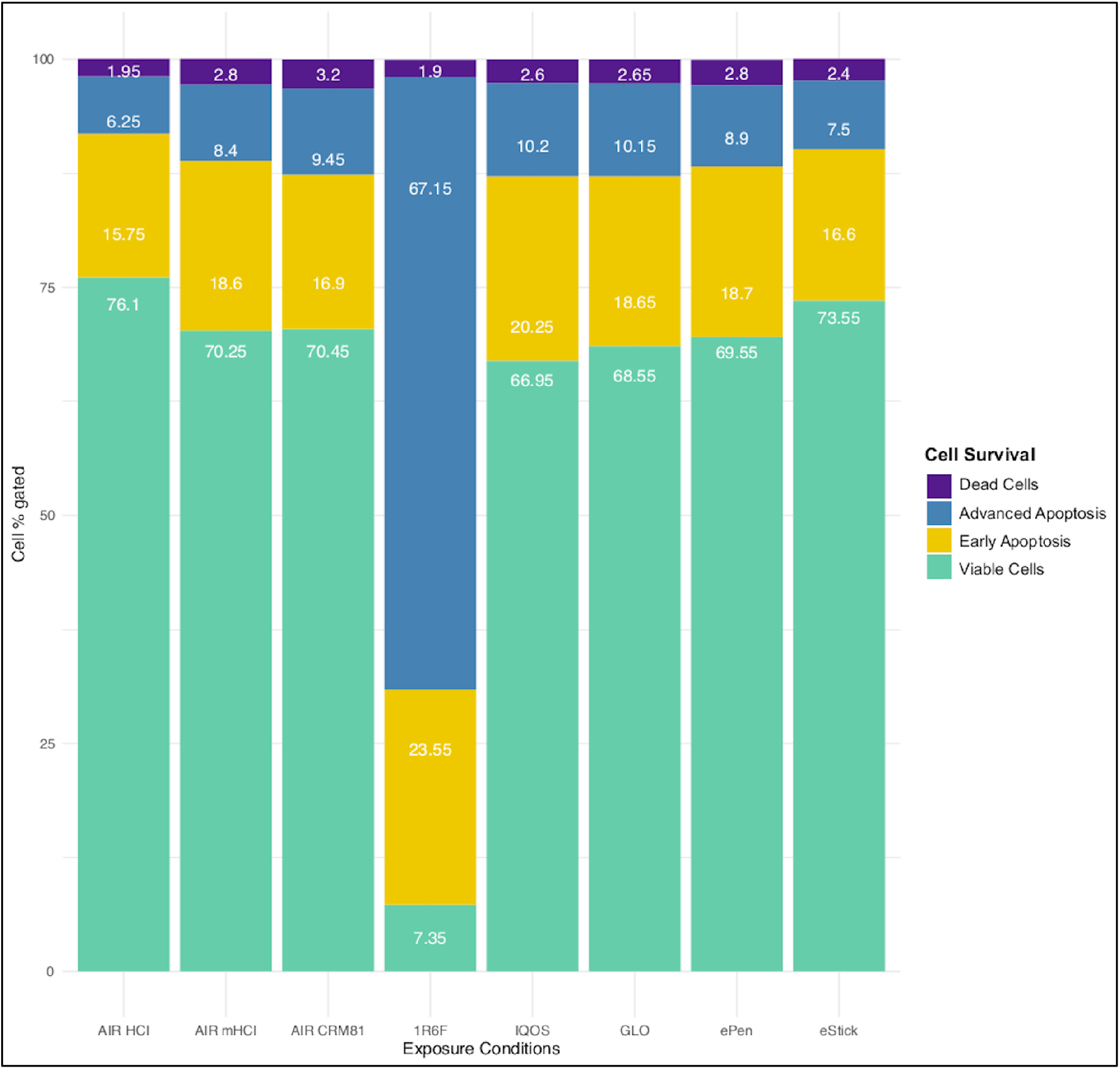
Evaluation of apoptosis and necrosis after smoke/vapor exposure. Each bar in the chart represents the whole (100%) of cell gated for the tested products, and segments in each bar represent the percentage of viable cells, early apoptotic cells, advanced apoptotic cells, and dead cells. Data were reported as percentage of cell gated.

### 3.4 Cell viability by NRU assay in live imaging

An alternative approach of NRU assay was performed using the Operetta High-Content Imaging System (PerkinElmer). H292 cells labelled with Neutral Red dye were monitored for 48 h after exposures (Figure 5). MFI values were normalised with the number of viable cells (labelled with NUCblue dye), and cell viability was expressed as MFI percentage of related air control. We observed a significant difference among the H292 cell viability curves during the observation time (p< 0.0001). 1R6F cell viability curve was much lower compared to AIR control and other product curves, whereas the trends of NGPs were more similar to each other with a cell proliferation % reduction higher than 1R6F. Particularly, exposure to 1R6F smoke significantly decreased H292 cell viability compared to AIR-HCI control starting from the 2^th^ hour (p< 0.0001) until 24 h (p< 0.0001) and 48 h (p< 0.0001). Moreover, H292 cell viability curve of 1R6F was significantly reduced compared to all the other tested products at 24 h (1R6F *vs* IQOS p value= 0.02; 1R6F *vs* GLO p value< 0.001; 1R6F *vs* ePen p value< 0.001; 1R6F *vs* eStick p value= 0.008). However, no significant decrease of H292 cell viability curve of 1R6F compared to GLO, ePen and eStick were observed at 48 h (p values> 0.05). Instead, the H292 cell viability curve of IQOS was not different compared to 1R6F curve until 11 h, but it was significant different from 12h to 48 h (p values> 0.05).

**Figure 5.**
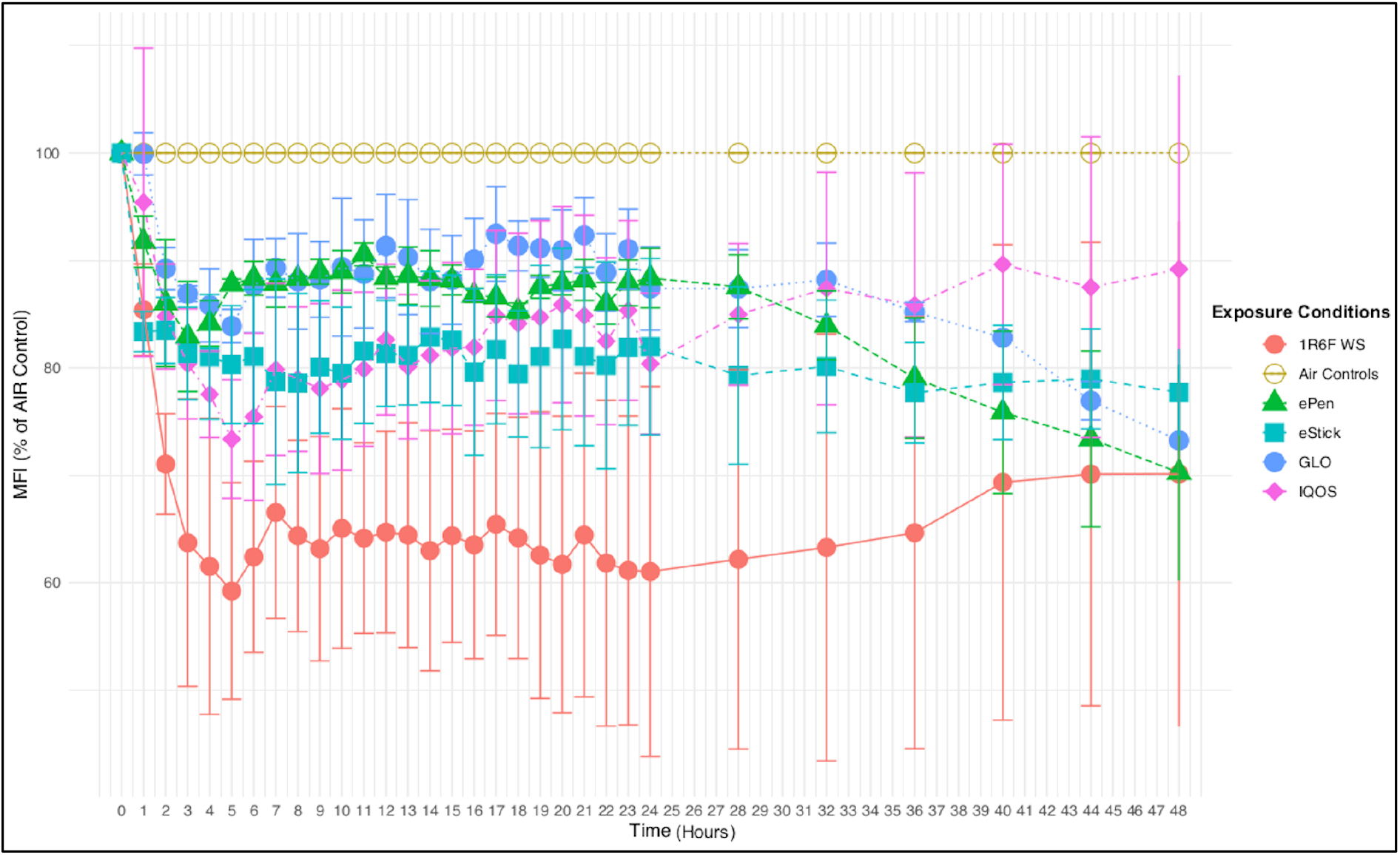
Cell viability by NRU using Operetta live imaging system. Data are reported as MFI percentage of related AIR control and showed as mean±SD.

Furthermore, the comparison between GLO curve and AIR-HCI was significantly different at 24 h (p= 0.017), but it was not significant at 48h. No significant differences were observed in H292 cell viability curves of GLO compared to AIR-HCI control at 24h, but this comparison becomes significant at 48 h (p<0.001). Also, the comparison between H292 cell viability curve of ePen and AIR-mHCI control was not significant at 24h, but it was significant at 48h (p< 0.0001). Instead, H292 cell viability curve of eStick was significantly different compared to AIR-CRM81 control both at 24 h (p= 0.038) and 48h (p= 0.004). The comparison among the H292 cell viability curves of AIR controls showed no significant differences. Linear regression analysis showed a significant relationship among the AIR controls (p< 0.0001). Additionally, significant relationship was observed among 1R6F, IQOS and GLO *vs* AIR-HCI control (p< 0.0001). Significant associations were also observed for ePen *vs* AIR-mHCI (p< 0.0001) control and eStick *vs* AIR-CRM81 control (p< 0.0001).

### 3.5 Real-Time Analysis of toxicity

Real-Time Cell viability analysis of H292 cell exposed to 1R6F cigarette smoke, THPs (IQOS and GLO), and e-cig (ePen and eStick) was also evaluated immediately after the exposures for 48 h (Figure 6). Cell index values were normalised with mean control value and data were expressed as percentages of related air control values. A significant difference among the H292 cell viability curves was observed over time (p< 0.0001). As expected, 1R6F cell viability curve was much lower compared to AIR control with significant differences for all the analysed time points (p< 0.0001). THPs cell viability curves showed a similar trend with significant decreases from AIR control both at 24 h (IQOS p< 0.0001; GLO p= 0.003) and 48 h (IQOS p< 0.0001; GLO p< 0.001). However, no significant differences were shown between THPs and 1R6F cell viability curves. Moreover, THPs cell viability curves were significantly lower compared to eStick both at 24 h (IQOS p< 0.0001; GLO p< 0.0001) and 48 h (IQOS p< 0.0001; GLO p< 0.0001), but no significant differences were showed between THPs and ePen cell viability curves. H292 cell viability curve after exposure to ePen vapor was not different from AIR control at 24h (p> 0.05), but a significant decrease was observed at 48 h (p= 0.01). Furthermore, ePen cell viability was significantly increased compared to 1R6F at 24h (p=0.002) but no difference was observed at 48 h (p>0.05). Exposure to eStick vapor showed a very different cell viability trend compared to AIR control and the other products. Indeed, cell viability of eStick was significantly increased already after 1h compared to both AIR control (p< 0.0001) and the other products (p< 0.0001). eStick cell viability started to decrease after approximately 24 h until 48 h, but, in spite of this, the differences remained significant compared to AIR control (p<0.001) and to all the other products (p< 0.0001).

**Figure 6.**
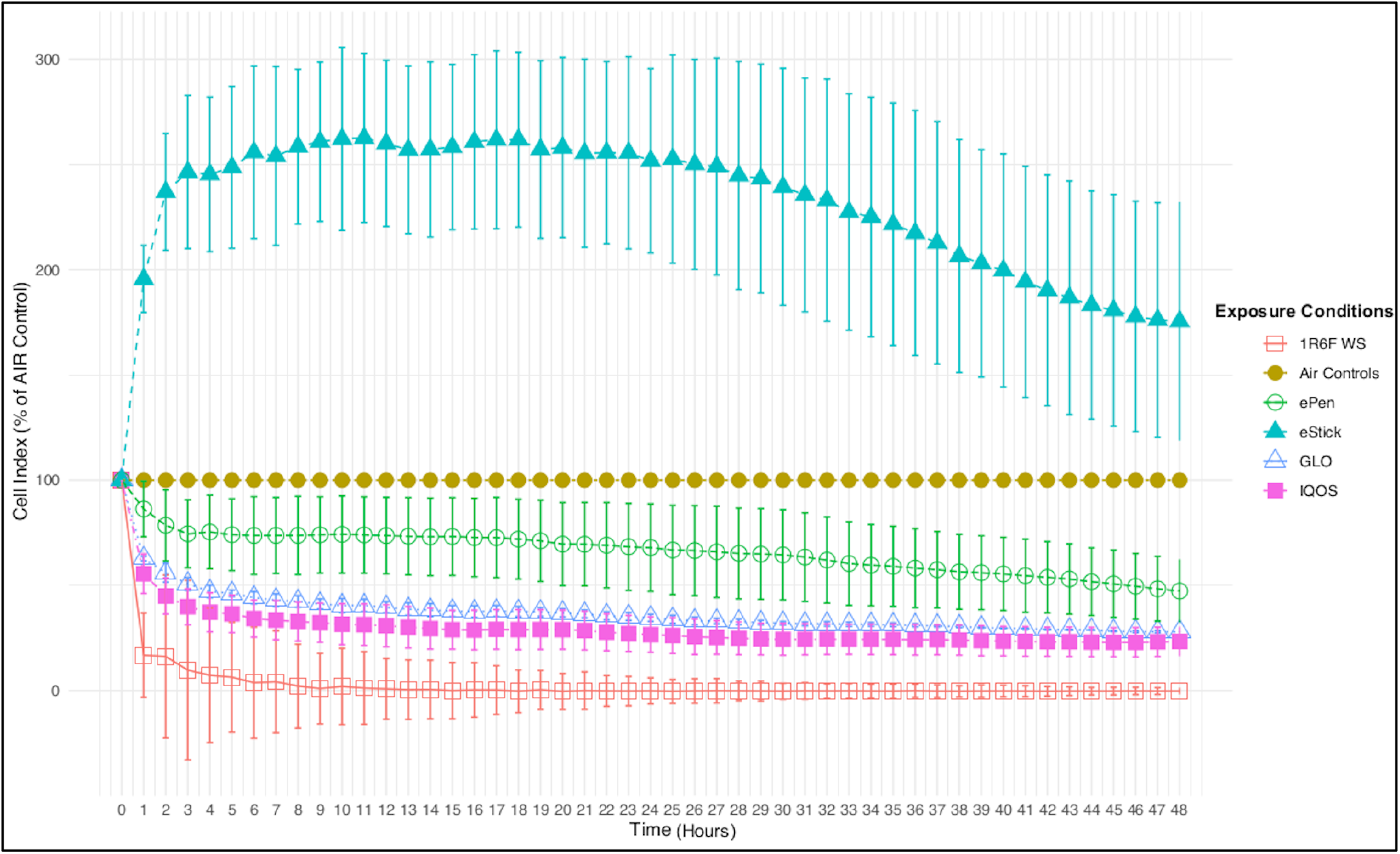
xCELLigence Real-Time Cell viability Analysis. Data are reported as Cell Index percentage of related AIR control and showed as mean±SD.

### 3.6 Comparison of cytotoxicity assays

Percentages of viable cells observed with the different cytotoxicity assays are reported in table 1. Cross-comparison results are shown in Tables A1 (1R6F), A2 (ePen), A3 (eStick), A4 (IQOS), and A5 (GLO) of the Appendix A. Even though all the cytotoxicity assays showed decreased cell viability after 1R6F exposure, we observed significant differences among them (p< 0.001). Indeed, almost all of cross-comparisons were significant (p values< 0.05). Particularly, substantial reduction in cell viability after 1R6F exposure was observed with Annexin V and RTCA assays compared with other assays (p values < 0.01). Also, the comparisons among NRU, MTT, Annexin V, NRU HCS, and RTCA assays were significantly different. But, the cross-comparisons showed that only the comparisons with RTCA assay were significantly different (p values < 0.05).

**Table 1.** Comparison of viable cells percentages obtained with different cytotoxicity assays at 24 hours after exposure.

|  | **NRU** | **MTT** | **Annexin V** | **NRU HCS** | **RTCA** | **P value** |
| --- | --- | --- | --- | --- | --- | --- |
| **1R6F** | 32.029 ± 1.78% | 49.21 ± 9.95% | 9.66 ± 1.58 % | 61.03 ± 17.22% | -0.1 ± 6.17% | <0.001 |
| **IQOS** | 88.315 ± 3.88% | 97.29 ± 9.95 % | 87.98 ± 8.83% | 80.38 ± 6.57% | 26.64 ± 8.41% | <0.001 |
| **GLO** | 89.984 ± 3.99% | 96.79 ± 8.54 % | 90.08 ± 1.21% | 87.39 ± 6.57% | 34.58 ± 1.87% | <0.001 |
| **ePen** | 95.631 ± 3.97% | 90.08 ± 2.79 % | 99.00 ± 0.50% | 88.36 ± 2.76% | 67.96 ± 20.74% | 0.003 |
| **eStick** | 101.586 ± 4.06% | 86.69 ± 3.63 % | 104.4 ± 1.10% | 81.97 ± 8.21% | 251.74 ± 43.77% | <0.001 |
| All data are reported as mean ± SD of percentage of their respectively AIR control. P values were calculated using one-way ANOVA. |  |  |  |  |  |  |

## 4. Discussion

Tobacco smoke consists of a complex mixture of gaseous and particulate components as a result of the combustion of organic matter. The cytotoxic effects are particularly relevant on bronchial epithelial cells which are directly exposed to smoke in human lungs. Different approaches were used to test and quantify these effects (Li, 2016; Putnam et al., 2002). The marketing of ENDS like THP and e-cig during the last decade, requires inclusion of new experimental protocols for comparative assessment of tobacco harm reduction between burning cigarettes and no-burning products (Belushkin et al., 2014). These assays are important to inform the regulatory bodies about their safety profile. It is therefore necessary to compare the cytotoxicity induced by ENDS with that induced by tobacco cigarettes and to evaluate the possible reduction of the damage triggered by these novel devices. One of the most used methods to evaluate cytotoxicity induced by ENDS vapor vs cigarette smoke is NRU assay. A low cost and easy test for indirect measurement of cell viability. But this assay, as many other cell viability and toxicity tests, has its own limitations (The National Toxicology Program (NTP) Interagency Center for the Evaluation of Alternative Toxicological Methods (NICEATM), 2003). In fact, any chemical affecting lysosomes may alter the result of the cell count and viability. This factor makes the system particularly sensitive to chemicals capable of specifically altering the lysosomal pH and cellular environment with consequent overestimation of the toxic effect. Considering the acidifying capacity of cigarette smoke, the possibility of altering the result with a pH-sensitive dye cannot be excluded. Previous reports showed no significant differences between NRU and MTT cytotoxicity evaluations with cigarette smoke and THP vapor (Davis et al., 2019). However, different studies concluded that NRU assay was the most sensitive cytotoxicity assay for moderate (3-6 h) and longer (12, 18 and 24 h) exposure times to smoke condensate compared to other cytotoxicity assays such as LDH release, kenacid blue binding, MTT, XTT, acid phosphatase activity, sulforhodamine B binding and resazurin binding (Putnam et al., 2002). Unfortunately, these studies, exposed bronchial epithelial cells to condensate smoking extract (CSE), and thus only to the water-soluble components of smoke/vapor, and therefore cannot be considered as reference studies in the toxicological assessment of cigarettes and ENDS. However, they have raised an important technical issue regarding the gold standard for toxicological evaluations. Therefore, we compared the standard NRU assay with the MTT assay (i), Annexin V Apoptosis Assay (ii), with an updated version of the NRU assay combined with HCS analysis (iii) and, finally, with a RTCA technology (iv) to evaluate the performance of these tests in the assessment of cigarette smoke and ENDS vapor induced cytotoxicity in a whole smoke/vapor ALI-exposed model of bronchial epithelial cells (H292). Our results, independently from the test used, showed an overall reduced toxicity in bronchial epithelial cells exposed to ENDS vapor when compared to cigarette smoke exposure.

However, different cytotoxicity results were observed for each assay on the basis of the evaluated cytotoxic mechanism and the peculiarity of each test. In particular, NRU and MTT results showed that 5 puffs of smoke from 1R6F cigarette were sufficient to adversely affect the uptake of dye and the metabolism of H292 bronchial cells. We also observed a slight significant difference of viable cells with NRU assay and MTT assay (Table A1). The MTT assay detects a higher percentage of cells survived to the exposure to 1R6F cigarette smoke. On the other hand, different results were obtained from NRU and MTT assays for H292 cells exposed to vapor of e-cig and THPs (Table A2–A5). Both, MTT and NRU assays did not show any difference between exposures to e-cigarettes and their AIR controls (Table A2, A3). Similarly, no significant differences between exposures to THPs and AIR control were detected by MTT assay (Figure 3). Our results are consistent with previous studies showing similar results on reduced cytotoxicity of e-cig and THPs compared to cigarette smoke (Davis et al., 2019; Leigh et al., 2018; Sohal et al., 2019). We also observed a significant reduction of H292 cell viability between exposures to THPs and their AIR controls by NRU assay (Figure 2), suggesting that this assay is more sensitive compared to MTT for the cytotoxicity evaluation of THPs.

The reduction of MTT leads to the formation of insoluble violet-blue formazan product, which are proportional to cell viability. Therefore, the acidic pH of the medium can modify the absorption spectrum of the cationic formazan altering the outcome of this test and representing an important bias for the interpretation of the results, particularly when it comes to tobacco products known to modify the pH of the medium (Brunnemann & Hoffmann, 1974; Henningfield et al., 1999). This aspect has to be considered when interpreting data from MTT assay on the effect of a complex chemical mixture, such as tobacco smoke and ENDS vapor, on cells, both to correctly interpret the result obtained and to evaluate a possible overestimation of viable cells. The reduction of pH by cigarette smoke on the culture medium could explain the high viability detected by MTT assay compared to NRU assay with 1R6F cigarette smoke, differently from ENDS vapor exposure, particularly for e-cig vapor, which does not contain tobacco (Table 1).

Our results obtained by Annexin V evaluation showed high reduction of viable cells after exposure to 1R6F [Annexin V-PE (−) and 7AAD (−)] compared to NRU and MTT assays with a substantial percentage of cells in advanced apoptosis [Annexin V-PE (+) and 7AAD (+)]. Comparing the percentage of viable cells obtained by the Annexin V apoptosis assay to that obtained by MTT assay, the result of viable cells by the latter assay is even higher. MTT is mainly reduced by the coenzyme NAD(P)H and glycolytic enzymes of the endoplasmic reticulum (Berridge et al., 2005). Since in cells undergoing early apoptosis coenzyme NAD(P)H and glycolytic enzymes remain intact and able to reduce MTT to formazan, the formation of this colored product in MTT assay could be viewed as a measure of the rate of glycolytic NAD(P)H production. Hence, a decrease in the concentration of D-glucose, NADH or NADPH in the culture medium may be accompanied by a decrease in MTT-formazan production. Another reason why the results of viable cells by MTT assay are higher may be due to test product different pH levels affecting the culture medium, to which the NRU assay does not seem to be sensitive. Our results suggest that for each product the results obtained by Apoptosis Annexin V assay are quite similar in terms of viable cells obtained with the classical NRU assay (Figure 4 and 2) due to the integrity of cell membranes in the early phase of apoptosis. Finally, effects of chemical mixture contained in smoke/vapor could result in changes in enzymatic activity that could influence the results of NRU and MTT assays (Winikoff et al., 2005). Cytotoxicity analysis by all these assays have the limit to detect live/dead cells at a given time (24 hours), without the possibility of understanding how cells behave in a continuum. Therefore, we also performed an NRU imaging assay, preparing a protocol on live imaging for NRU in lysosomes, then observing the morphological changes and the phenotypic fingerprinting, indicator of metabolically active cells over the time (from 0 to 48 hours), going therefore beyond the determination of cytotoxicity in classical tests.

Finally, the RTCA data are produced and collected during the whole time of running protocol, and hence there is the possibility of assessing the effect of smoke/vapor on cells at any time of the post-exposure time (0 - 48 hours), generating proliferation profiles in one single experiment (Figure 6). By comparing of the results from this technique and the others we observed a strong deviation from the others at 24 hours, in particular for the 1R6F cigarette smoke and for THP vapor exposure. Despite a good number of viable cells for e-cigs, there is a significant difference between the data obtained for the other products. The fact that this method mainly detects the ability of cells to attach and proliferate suggest that cells are viable and metabolically active. However, our results suggest that cells exhibit delayed replication as suggested by low cellular indexes following treatment with cigarette smoke and THP vapor, but not with treatment with e-cig vapor (Figure 6 and Table 1). This data needs to be further investigated to better understand the limitations or benefits of RTCA for the assessment of these products. RTCA and HCS still have the advantage to generate a time-dependent growth/mortality profile in a single experimental run, differently from other assays. This is a relevant point allowing to determine and derive the kinetic dependency parameter for effectiveness and potency of smoke/vapor in the cell metabolism. However, these methods have some limitations related to the costs of the instruments and personnel training for non-routinary methodology. Increased costs for performing such tests are also depending on those related for buying dedicated disposable parts for imaging and RTCA.

## 5. Conclusion

In conclusion, MTT assay applied to the assessment of the cytotoxicity induced by cigarette smoke and ENDS vapors could suffer from potential interference due to the intrinsic nature of these chemical mixtures. Instead, NRU assay is a more appropriate test for assessing the cytotoxicity induced by these products. However, it should be combined with a time-resolved test, particularly when studying new products and devices whose kinetics are unknown. Furthermore, it would be advisable to deepen the results of the NRU assay with a flow cytometric test evaluating apoptosis and necrosis in order to increase toxicological sensitivity.

## Abbreviations

ENDS: Electronic Nicotine Delivery System
THPs: Tobacco Heating Products
E-cigs: Electronic Cigarettes
NRU: Neutral Red Uptake
NR: Neutral Red
MTT: 3-(4,5-dimethylthiazol-2-yl)-2,5-diphenyltetrazolium bromide
HCS: High Content Screening
RTCA: Real Time Cell Analysis
ISO: International Organization for Standardization
HCI: Health Canada Intensive
CRM81: CORESTA Recommended Method n. 81

## APPENDIX A.

**Table A1.** Cross-comparisons among NRU, MTT, Annexin V, NRU-HCS, and RTDA assessing the toxicity of 1R6F reference cigarette.

| **Adj p values**<br>**1R6F** | **NRU** | **MTT** | **Annexin V** | **NRU-HCS** | **RTCA** |
| --- | --- | --- | --- | --- | --- |
| **NRU** |  | 0.036* | 0.058 | 0.004** | 0.005** |
| **MTT** | 0.036* | | 0.0008<sup>#</sup> | 0.386 | <0.0001<sup>\$</sup> |
| **Annexin V** | 0.058 | 0.0008<sup>#</sup> |  | 0.0002<sup>#</sup> | 0.814 |
| **NRU-HCS** | 0.004** | 0.386 | 0.0002<sup>#</sup> | | <0.0001<sup>\$</sup> |
| **RTCA** | 0.005** | <0.0001<sup>\$</sup> | 0.814 | <0.0001<sup>\$</sup> | |
| Adjusted p values were calculated using Tukey's adjustment for multiple comparisons.<br>Significance code: < 0.0001 (\$), < 0.001 (#), < 0.01(**), < 0.05 (*) | | | | | |

**Table A2.** Cross-comparisons among NRU, MTT, Annexin V, NRU-HCS, and RTDA assessing the toxicity of ePen vapor.

| **Adj p values**<br>**ePen** | **NRU** | **MTT** | **Annexin V** | **NRU-HCS** | **RTCA** |
| --- | --- | --- | --- | --- | --- |
| **NRU** |  | 0.762 | 0.985 | 0.716 | 0.002** |
| **MTT** | 0.762 |  | 0.671 | 0.998 | 0.012* |
| **Annexin V** | 0.985 | 0.671 |  | 0.617 | 0.006** |
| **NRU-HCS** | 0.716 | 0.998 | 0.617 |  | 0.052 |
| **RTCA** | 0.002** | 0.012* | 0.006** | 0.052 |  |
| Adjusted p values were calculated using Tukey's adjustment for multiple comparisons.<br>Significance code: < 0.0001 (§), < 0.001 (#), < 0.01(**), < 0.05 (*) |  |  |  |  |  |

**Table A3.** Cross-comparisons among NRU, MTT, Annexin V, NRU-HCS, and RTDA assessing the toxicity of eStick vapor.

| **Adj p values**<br>**eStick** | **NRU** | **MTT** | **Annexin V** | **NRU-HCS** | **RTCA** |
| --- | --- | --- | --- | --- | --- |
| **NRU** |  | 0.999 | 0.999 | 0.331 | <0.0001<sup>§</sup> |
| **MTT** | 0.999 |  | 0.996 | 0.544 | <0.0001<sup>§</sup> |
| **Annexin V** | 0.999 | 0.996 |  | 0.439 | <0.0001<sup>§</sup> |
| **NRU-HCS** | 0.331 | 0.544 | 0.439 |  | <0.0001<sup>§</sup> |
| **RTCA** | <0.0001<sup>§</sup> | <0.0001<sup>§</sup> | <0.0001<sup>§</sup> | <0.0001<sup>§</sup> |  |
| Adjusted p values were calculated using Tukey's adjustment for multiple comparisons.<br>Significance code: < 0.0001 (§), < 0.001 (#), < 0.01(**), < 0.05 (*) |  |  |  |  |  |

**Table A4.** Cross-comparisons among NRU, MTT, Annexin V, NRU-HCS, and RTDA assessing the toxicity of IQOS.

| **Adj p values**<br>**IQOS** | **NRU** | **MTT** | **Annexin V** | **NRU-HCS** | **RTCA** |
| --- | --- | --- | --- | --- | --- |
| **NRU** |  | 0.227 | 1 | 0.52 | <0.0001<sup>§</sup> |
| **MTT** | 0.227 |  | 0.505 | 0.027* | <0.0001<sup>§</sup> |
| **Annexin V** | 1 | 0.505 |  | 0.759 | <0.0001<sup>§</sup> |
| **NRU-HCS** | 0.52 | 0.027* | 0.759 |  | <0.0001<sup>§</sup> |
| **RTCA** | <0.0001<sup>§</sup> | <0.0001<sup>§</sup> | <0.0001<sup>§</sup> | <0.0001<sup>§</sup> |  |
| Adjusted p values were calculated using Tukey's adjustment for multiple comparisons.<br>Significance level: < 0.0001 (§), < 0.001 (#), < 0.01(**), < 0.05 (*) |  |  |  |  |  |

**Table A5.** Cross-comparisons among NRU, MTT, Annexin V, NRU-HCS, and RTDA assessing the toxicity of GLO.

| **Adj p values**<br>**GLO** | **NRU** | **MTT** | **Annexin V** | **NRU-HCS** | **RTCA** |
| --- | --- | --- | --- | --- | --- |
| **NRU** | | 0.308 | 1 | 0.968 | <0.0001<sup>\$</sup> |
| **MTT** | 0.308 | | 0.634 | 0.21 | <0.0001<sup>\$</sup> |
| **Annexin V** | 1 | 0.634 | | 0.986 | <0.0001<sup>\$</sup> |
| **NRU-HCS** | 0.968 | 0.21 | 0.986 | | <0.0001<sup>\$</sup> |
| **RTCA** | <0.0001<sup>\$</sup> | <0.0001<sup>\$</sup> | <0.0001<sup>\$</sup> | <0.0001<sup>\$</sup> | |
| Adjusted p values were calculated using Tukey's adjustment for multiple comparisons.<br>Significance level: < 0.0001 (\$), < 0.001 (#), < 0.01(**), < 0.05 (*) | | | | | |

## References

Azzopardi, D., Haswell, L. E., Foss-Smith, G., Hewitt, K., Asquith, N., Corke, S., & Phillips, G. (2015, Oct). Evaluation of an air-liquid interface cell culture model for studies on the inflammatory and cytotoxic responses to tobacco smoke aerosols. Toxicol In Vitro, 29(7), 1720–1728. https://doi.org/10.1016/j.tiv.2015.06.016

Azzopardi, D., Patel, K., Jaunky, T., Santopietro, S., Camacho, O. M., McAughey, J., & Gaça, M. (2016, Jul). Electronic cigarette aerosol induces significantly less cytotoxicity than tobacco smoke. Toxicol Mech Methods, 26(6), 477–491. https://doi.org/10.1080/15376516.2016.1217112

Baker, R. R., Pereira da Silva, J. R., & Smith, G. (2004a). The effect of tobacco ingredients on smoke chemistry. Part I: Flavourings and additives. Food Chem Toxicol, 42 Suppl, S3–37. https://doi.org/10.1016/s0278-6915(03)00189-3

Baker, R. R., Pereira da Silva, J. R., & Smith, G. (2004b). The effect of tobacco ingredients on smoke chemistry. Part II: casing ingredients. Food Chem Toxicol, 42 Suppl, S39–52. https://doi.org/10.1016/j.fct.2003.08.009

Bekki, K., Inaba, Y., Uchiyama, S., & Kunugita, N. (2017). Comparison of Chemicals in Mainstream Smoke in Heat-not-burn Tobacco and Combustion Cigarettes. J uoeh, 39(3), 201–207. https://doi.org/10.7888/juoeh.39.201

Belushkin, M., Piadé, J. J., Chapman, S., & Fazekas, G. (2014, Mar). Investigating predictability of in vitro toxicological assessments of cigarettes: analysis of 7 years of regulatory submissions to Canadian regulatory authorities. Regul Toxicol Pharmacol, 68(2), 222–230. https://doi.org/10.1016/j.yrtph.2013.12.009

Berridge, M. V., Herst, P. M., & Tan, A. S. (2005). Tetrazolium dyes as tools in cell biology: new insights into their cellular reduction. Biotechnol Annu Rev, 11, 127–152. https://doi.org/10.1016/s1387-2656(05)11004-7

Brunnemann, K. D., & Hoffmann, D. (1974, Feb). The pH of tobacco smoke. Food Cosmet Toxicol, 12(1), 115–124. https://doi.org/10.1016/0015-6264(74)90327-7

Center for Tobacco Reference Products Kentucky University. (2018). Certificate of Analysis: 1R6F Certified Reference Cigarette. https://ctrp.uky.edu/assets/pdf/webdocs/CoA18_1R6F.pdf

Comer, D. M., Kidney, J. C., Ennis, M., & Elborn, J. S. (2013, May). Airway epithelial cell apoptosis and inflammation in COPD, smokers and nonsmokers. Eur Respir J, 41(5), 1058–1067. https://doi.org/10.1183/09031936.00063112

CORESTA. (2004). The CORESTA E-Cigarette Task Force. The rationale and strategy for conducting in vitro toxicology testing of tobacco smoke. https://www.coresta.org/sites/default/files/technical_documents/main/IVT_TF_Rationale-IVT-Testing-Tob.-Smoke_Report_Jun04.pdf

Davis, B., Dang, M., Kim, J., & Talbot, P. (2015, Feb). Nicotine concentrations in electronic cigarette refill and do-it-yourself fluids. Nicotine Tob Res, 17(2), 134–141. https://doi.org/10.1093/ntr/ntu080

Davis, B., To, V., & Talbot, P. (2019, Dec). Comparison of cytotoxicity of IQOS aerosols to smoke from Marlboro Red and 3R4F reference cigarettes. Toxicol In Vitro, 61, 104652. https://doi.org/10.1016/j.tiv.2019.104652

Dempsey, R., Coggins, C. R., & Roemer, E. (2011, Oct). Toxicological assessment of cigarette ingredients. Regul Toxicol Pharmacol, 61(1), 119–128. https://doi.org/10.1016/j.yrtph.2011.07.002

Eaton, D., Jakaj, B., Forster, M., Nicol, J., Mavropoulou, E., Scott, K., Liu, C., McAdam, K., Murphy, J., & Proctor, C. J. (2018, 2018/03/01/). Assessment of tobacco heating product THP1.0. Part 2: Product design, operation and thermophysical characterisation. Regulatory Toxicology and Pharmacology, 93, 4–13. https://doi.org/https://doi.org/10.1016/j.yrtph.2017.09.009

European Commission. (2000). Commission Directive 2000/33/EC of 25 April 2000 adapting to technical progress for the 27th time Council Directive 67/548/EEC on the approximation of laws, regulations and administrative provisions relating to the classification, packaging and labelling of dangerous substances (Text with EEA relevance.), CELEX1. Publications Office of the European Union. http://op.europa.eu/en/publication-detail/-/publication/eeb87812-0c0b-49f1-a0fd-edebccb68ce5

Henningfield, J. E., Fant, R. V., Radzius, A., & Frost, S. (1999, Jun). Nicotine concentration, smoke pH and whole tobacco aqueous pH of some cigar brands and types popular in the United States. Nicotine Tob Res, 1(2), 163–168. https://doi.org/10.1080/14622299050011271

Imai, K., Mercer, B. A., Schulman, L. L., Sonett, J. R., & D’Armiento, J. M. (2005, Feb). Correlation of lung surface area to apoptosis and proliferation in human emphysema. Eur Respir J, 25(2), 250–258. https://doi.org/10.1183/09031936.05.00023704

Iskandar, A. R., Gonzalez-Suarez, I., Majeed, S., Marescotti, D., Sewer, A., Xiang, Y., Leroy, P., Guedj, E., Mathis, C., Schaller, J. P., Vanscheeuwijck, P., Frentzel, S., Martin, F., Ivanov, N. V., Peitsch, M. C., & Hoeng, J. (2016, Jul). A framework for in vitro systems toxicology assessment of e-liquids. Toxicol Mech Methods, 26(6), 389–413. https://doi.org/10.3109/15376516.2016.1170251

Johnson, M. D., Schilz, J., Djordjevic, M. V., Rice, J. R., & Shields, P. G. (2009, Dec). Evaluation of in vitro assays for assessing the toxicity of cigarette smoke and smokeless tobacco. Cancer Epidemiol Biomarkers Prev, 18(12), 3263–3304. https://doi.org/10.1158/1055-9965.Epi-09-0965

Kim, S., Goniewicz, M. L., Yu, S., Kim, B., & Gupta, R. (2015, May 5). Variations in label information and nicotine levels in electronic cigarette refill liquids in South Korea: regulation challenges. Int J Environ Res Public Health, 12(5), 4859–4868. https://doi.org/10.3390/ijerph120504859

Leigh, N. J., Tran, P. L., O’Connor, R. J., & Goniewicz, M. L. (2018, Nov). Cytotoxic effects of heated tobacco products (HTP) on human bronchial epithelial cells. Tob Control, 27(Suppl 1), s26–s29. https://doi.org/10.1136/tobaccocontrol-2018-054317

Li, X. (2016, Oct). In vitro toxicity testing of cigarette smoke based on the air-liquid interface exposure: A review. Toxicol In Vitro, 36, 105–113. https://doi.org/10.1016/j.tiv.2016.07.019

Li, X., Luo, Y., Jiang, X., Zhang, H., Zhu, F., Hu, S., Hou, H., Hu, Q., & Pang, Y. (2019, Jan 1). Chemical Analysis and Simulated Pyrolysis of Tobacco Heating System 2.2 Compared to Conventional Cigarettes. Nicotine Tob Res, 21(1), 111–118. https://doi.org/10.1093/ntr/nty005

Mandavilli, B. S., Aggeler, R. J., & Chambers, K. M. (2018). Tools to Measure Cell Health and Cytotoxicity Using High Content Imaging and Analysis. Methods Mol Biol, 1683, 33–46. https://doi.org/10.1007/978-1-4939-7357-6_3

OECD/OCDE. (2004). OECD GUIDELINE FOR TESTING OF CHEMICALSGuideline - 432 in vitro 3T3 nru phototoxicity test. OECD Better Policies for Better Lives. http://www.oecd.org/env/ehs/testing/TG432-TC-4dec-clean.pdf

Putnam, K. P., Bombick, D. W., & Doolittle, D. J. (2002, Oct). Evaluation of eight in vitro assays for assessing the cytotoxicity of cigarette smoke condensate. Toxicol In Vitro, 16(5), 599–607. https://doi.org/10.1016/s0887-2333(02)00050-4

Repetto, G., del Peso, A., & Zurita, J. L. (2008). Neutral red uptake assay for the estimation of cell viability/cytotoxicity. Nat Protoc, 3(7), 1125–1131. https://doi.org/10.1038/nprot.2008.75

Sohal, S. S., Eapen, M. S., Naidu, V. G. M., & Sharma, P. (2019, Feb). IQOS exposure impairs human airway cell homeostasis: direct comparison with traditional cigarette and e-cigarette. ERJ Open Res, 5(1). https://doi.org/10.1183/23120541.00159-2018

Stockert, J. C., Horobin, R. W., Colombo, L. L., & Blázquez-Castro, A. (2018, Apr). Tetrazolium salts and formazan products in Cell Biology: Viability assessment, fluorescence imaging, and labeling perspectives. Acta Histochem, 120(3), 159–167. https://doi.org/10.1016/j.acthis.2018.02.005

The National Toxicology Program (NTP) Interagency Center for the Evaluation of Alternative Toxicological Methods (NICEATM). (2003). A Test for Basal Cytotoxicity for an In Vitro Validation Study Phase III (National Institute of Environmental Health Sciences (NIEHS)-National Institutes of Health (NIH) U.S. Public Health Service - Department of Health and Human Services, Issue. https://www.google.com/url?sa=t&rct=j&q=&esrc=s&source=web&cd=&ved=2ahUKEwiEqfH4kf3uAhXIwAIHHRmXAPMQFjAAegQIARAD&url=https%3A%2F%2Fntp.niehs.nih.gov%2Ficcvam%2Fmethods%2Facutetox%2Finvidocs%2Fphiiiprot%2Fnhkphiii.pdf&usg=AOvVaw3RjJWwE10595gXu06E8IN-

Winikoff, S. E., Zeh, H. J., DeMarco, R., & Lotze, M. T. (2005). Chapter 29 - Cytolytic Assays. In M. T. Lotze & A. W. Thomson (Eds.), Measuring Immunity (pp. 343–349). Academic Press. https://doi.org/https://doi.org/10.1016/B978-012455900-4/50291-9

Yan, G., Du, Q., Wei, X., Miozzi, J., Kang, C., Wang, J., Han, X., Pan, J., Xie, H., Chen, J., & Zhang, W. (2018, Dec 11). Application of Real-Time Cell Electronic Analysis System in Modern Pharmaceutical Evaluation and Analysis. Molecules, 23(12). https://doi.org/10.3390/molecules23123280

